# Estimation of inbreeding and kinship coefficients via latent identity-by-descent states

**DOI:** 10.1101/2023.09.02.556031

**Authors:** Yongtao Guan, Daniel Levy

## Abstract

Estimating the individual inbreeding coefficient and pairwise kinship is an important problem in human genetics (e.g., in disease mapping) and in animal and plant genetics (e.g., inbreeding design). Existing methods such as sample correlation-based genetic relationship matrix, KING, and UKin are either biased, or not able to estimate inbreeding coefficients, or produce a large proportion of negative estimates that are difficult to interpret. This limitation of existing methods is partly due to failure to explicitly model inbreeding. Since all humans are inbred to various degrees by virtue of shared ancestries, it is prudent to account for inbreeding when inferring kinship between individuals. We present “Kindred”, an approach that estimates inbreeding and kinship by modeling latent identity-by-descent states that accounts for all possible allele sharing – including inbreeding – between two individuals. Through simulation, we demonstrate the high accuracy and, more importantly, non-negativity of kinship estimates by Kindred. By selecting a subset of SNPs that are similar in allele frequencies across different populations, Kindred can accurately estimate kinship between admixed samples. Finally, we demonstrate that the realized kinship matrix estimated by Kindred is effective in reducing genomic control values via linear mixed model in genome-wide association studies, and it also produces sensible heritability estimates. Kindred is freely available at http://www.haplotype.org.

## 1 Introduction

Kinship (denoted by *ϕ*) between two individuals is the probability that two alleles sampled at a random locus each from one individual are identical by descent (IBD). The inbreeding coefficient (denoted by *F*) of an individual is the probability that two parental alleles sampled at a random locus in the genome are IBD. Thus inbreeding coefficient of an individual is the kinship between his or her parents. In addition, between one and oneself (or between monozygotic twins) *ϕ* = (1 + *F*)*/*2 (Wright, 1922). Therefore, inbreeding can be treated as a derived concept of kinship, and a statistical model designed to estimate kinship automatically applies to estimating the inbreeding coefficient as well. Kinship estimated from genotype data is called realized kinship. Due to the stochastic nature of recombination and gamete segregation, realized kinship may have significant variation from pedigree estimates (Visscher et al., 2006). Estimating the individual inbreeding coefficient and pairwise kinship is an important problem in human disease mapping, forensics, animal and plant breeding, and conservation and evolutionary biology (Wang, 2016).

In an effort to study gene IBD, Jacquard (1972) documented nine IBD states between any two individuals within a pedigree. These IBD states are partially observable within a pedigree, and the mean probabilities of each state can be computed purely based on the pedigree. These IBD states, however, are completely latent between two individuals not linked by a known pedigree. Thompson (2013) described connections between the Jacquard IBD states and Ewen’s sampling partition in coalescence, and provided the joint distribution of genotypes between two individuals conditioning on latent gene IBD states. These formed the basis for our strategy to estimate kinship by inferring latent IBD states between two individuals via joint genotype distributions. A similar approach was implemented by Milligan (2003) using maximum likelihood, albeit only using a few dozen markers.

Existing methods of kinship estimation, such as KING (Manichaikul et al., 2010) and the recent UKin approach (Jiang et al., 2022) ignore the possible inbreeding of each sample. Inbreeding can affect the abundance of homozygous markers in the genome; failure to account for inbreeding results not only in the inability to estimate inbreeding, but also biases kinship estimates (more in the Discussion). The sample correlation-based genetic relatedness matrix (scGRM) has been widely used as the kinship matrix (Price et al., 2006; Yang et al., 2011; Wang et al., 2017). But it has been shown that scGRM estimates are biased downward (Weir and Goudet, 2017; Ochoa and Storey, 2021; Jiang et al., 2022), and this bias is due to scGRM using sample allele frequency in computation, which are evidently biased compare to the allele frequencies of the reference population. Several methods have been developed to correct for the bias. In particular, Weir and Goudet (2017) used least related samples to recalibrate the kinship estimates; Jiang et al. (2022) quantified the bias under an assumption of no inbreeding and correct the bias. But both approaches still produce negative estimates for unrelated samples. Ochoa and Storey (2021) achieved non-negativity by subtracting the smallest estimates; their method also ignored inbreeding by only considering locally outbred individuals.

We present Kindred, which estimates kinship and inbreeding coefficients by inferring the loading probabilities of latent IBD states. Although Kindred is a model-based method, its computation is efficient, owning to the least square approach we take. Through simulations, we demonstrate its high accuracy and non-negativity in kinship estimates. In addition, by selecting a subset of SNPs that are similar in allele frequencies across different populations, we demonstrate that Kindred accurately estimates kinship between admixed samples. A major application of the kinship matrix in genome-wide association studies is the variance-covariance matrix for random effect in linear mixed models, and we demonstrate that realized kinship matrix estimated by Kindred is effective in reducing genomic control values. Another application of the kinship matrix is to estimate heritability through variance component models, and we demonstrate that the realized kinship matrix estimated by Kindred produced a higher, statistically significant, heritability estimate than other methods.

## 2 Materials and Methods

### 2.1 Latent IBD states model

At an arbitrary marker, there are 15 detailed IBD states between four alleles of two individuals. If the parental origin of an allele is not of interest, these 15 states can be reduced to nine condensed IBD states (Jacquard, 1972; Thompson, 2013). Here we work with the nine condensed identity states detailed in Table 1. Conditional on each state, the distribution of joint genotypes for a bi-allelic SNP is given in Table 1. This distribution is a bi-allelic special case of what is presented for the four allelic case (Thompson, 2013). Kinship can be computed from the loading probabilities, ∆_*j*_ for *j*-th latent IBD state Σ_*j*_, as follows (Jacquard, 1972):

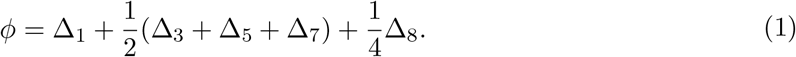

With reference to the diagram in Table 1, all latent states that have IBD between two samples (yellow segments) contribute to the kinship calculation in Equation 1. The fractional coefficients multiplying each loading probabilities in Equation 1 come from their respective numbers of equivalent and mutually exclusive IBD states. (Combining equivalent and mutually exclusive states is how 15 states were condensed to nine.) Inbreeding coefficients can also be computed from the loading probabilities of latent IBD states (Jacquard, 1972):

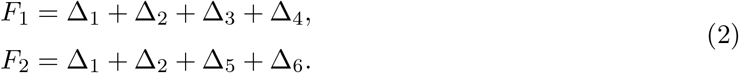

With Table 1, we can model the observed genotype as emanating from the mixture of the nine latent IBD states to estimate the loading probabilities (∆’s), and from which to estimate kinship using Equation (1). We start by considering a subset of SNPs with allele frequency *p* so that they share the same matrix in Table 1. Denote *n*_*p*,1_ the count of *AA AA, n*_*p*,2_ the count of *AA AB*, …, and *n*_*p*,9_ the count of *BB BB* (Table 1), the multinomial likelihood is

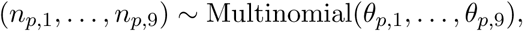

and the maximum likelihood estimate for 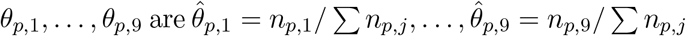. We can then model joint genotypes distribution 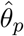 as a linear combination of Σ_*j*_’s to seek the least squares fit of a constrained system

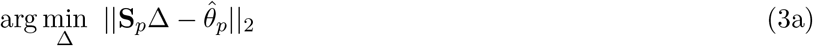

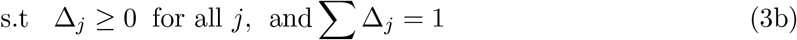

where **S**_*p*_ = (Σ_1_, …, Σ_9_) is a 9 *×* 9 matrix detailed in Table 1, ∆ = (∆_1_, …, ∆_9_) is the vector of loading probabilities, and || *·* ||_2_ denotes the *L*_2_ norm, which is the square root of sum of squares of all components. From the estimates of the loading probabilities 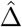, we can compute the kinship estimate 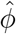 using Equation (1). In essence we treat 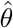 as observed and seek a non-negative least squares fit to obtain 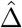 .

**Table 1:** The diagram above the table shows nine states of IBD sharing between two individuals. These are colored re-rendering of Table 2 in (Jacquard, 1972). Two alleles of individual 1 are colored in blue and two alleles of individual 2 in black. Two alleles IBD within an individual are connected via a purple segment. Two alleles IBD between individuals are connected by an orange segment. Σ_7_, Σ_8_, and Σ_9_ are two non-inbreed individuals share two, one, and zero alleles IBD respectively. The table details joint genotype distribution for each latent IBD states, reproduced from Table 3 of (Thompson, 2013). Each of the nine columns labelled by Σ_*j*_ corresponds to a latent state in the diagram above. The joint genotypes are listed in columns G1 and G2 where the order matters. *P* denotes the frequency of A allele and *q* frequency of B allele, and *p* + *q* = 1. Note each column labelled by Σ_*j*_ sums to 1. When G1 and G2 are the genotypes of the same individual, only Σ_1_ and Σ_7_ are relevant. When inbreedings are ignored, only last three Σ columns are relevant.

| $\Sigma_1$ | $\Sigma_2$ | $\Sigma_3$ | $\Sigma_4$ | $\Sigma_5$ | $\Sigma_6$ | $\Sigma_7$ | $\Sigma_8$ | $\Sigma_9$ | G1 | G2 |
| --- | --- | --- | --- | --- | --- | --- | --- | --- | --- | --- |
| $p$ | $p^2$ | $p^2$ | $p^3$ | $p^2$ | $p^3$ | $p^2$ | $p^3$ | $p^4$ | AA | AA |
| 0 | 0 | $pq$ | $2p^2q$ | 0 | 0 | 0 | $p^2q$ | $2p^3q$ | AA | AB |
| 0 | $pq$ | 0 | $pq^2$ | 0 | $p^2q$ | 0 | 0 | $p^2q^2$ | AA | BB |
| 0 | 0 | 0 | 0 | $pq$ | $2p^2q$ | 0 | $p^2q$ | $2p^3q$ | AB | AA |
| 0 | 0 | 0 | 0 | 0 | 0 | $2pq$ | $pq$ | $4p^2q^2$ | AB | AB |
| 0 | 0 | 0 | 0 | $pq$ | $2pq^2$ | 0 | $pq^2$ | $2pq^3$ | AB | BB |
| 0 | $pq$ | 0 | $p^2q$ | 0 | $pq^2$ | 0 | 0 | $p^2q^2$ | BB | AA |
| 0 | 0 | $pq$ | $2pq^2$ | 0 | 0 | 0 | $pq^2$ | $2pq^3$ | BB | AB |
| $q$ | $q^2$ | $q^2$ | $q^3$ | $q^2$ | $q^3$ | $q^2$ | $q^3$ | $q^4$ | BB | BB |

For the *i*-th SNP with allele frequency *p*_*i*_, we can compute 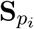 and we observe 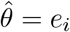, where *e*_*i*_ has a single entry equals 1 (depending on the joint genotypes) and the rest 8 entries 0. We append 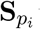’s together to obtain the design matrix **S** and append *e*_*i*_’s together to obtain a vector 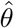. For total *m* SNPs, **S** is an 9*m ×* 9 matrix, and 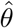 is an 9*m* vector, and we seek the least squares fit of a constrained system

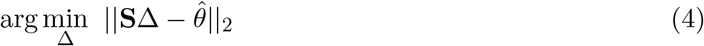

with constraints (3b). The above system generalizes (3) to SNPs of arbitrary allele frequencies. If all *p*_*i*_ = *p*, the above system reduces to (3) (below). In practice, we find binning SNPs and re-estimate allele frequencies in each bin improves performance (below).

### 2.2 Least squares fit

To fit a least squares arg min_*x*_ ||*Ax−b*||_2_ under constraints is equivalent to fit arg min_*x*_ ||*A*^*t*^*Ax−A*^*t*^*b*||_2_ under the same constraints. For a set of SNPs with the same allele frequency *p*, we have two ways to formulate *A* and *b*. One way is such that *A* = **S**_*p*_ and 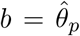, the other is 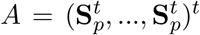 and 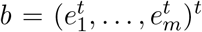. In the second formulation, 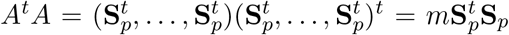 and 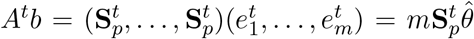, which is same as the first formulation multiplying *m* on both terms. Generalizing this to arbitrary allele frequencies, we can efficiently compute 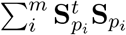 and 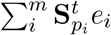, by rounding *p*_*i*_ to, say, the second decimal place to reuse the matrix 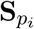 and fit an equivalent system arg min_∆_ 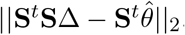.

### 2.3 Binning SNPs and the optimal bin size

We first round SNP allele frequencies to the second digit (i.e. with increment of 0.01), which we call rounded allele frequencies (RAF). For each RAF *f*, we bin SNPs in the range of *S*_*f*_ = (*f −b, f* + *b*), and for each pair of samples, we computed allele frequency using all SNPs in set *S*_*f*_ to obtain 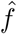, which was used to obtain 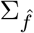. For most bins the differences between 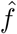 and *f* are small, but 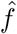 produced better estimates in simulations, presumably by alleviating the effects of mis-specification of allele frequencies. Binning SNPs gives us a tunable parameter *b*. To determine the optimal *b* we resort to simulations. Briefly, we simulate first cousins whose expected kinship is 0.0625, then we ran Kindred to estimate kinship coefficients with different *b* ranging from 0 to 0.10 with step size of 0.01. Supplementary Figure S1 plots the deviation from the truth for different bin sizes, and *b* = 0.04 appears to be optimal in all three sets of simulations using three continental population as founders.

## 3 Results

### 3.1 Kinship between non-admixed samples

We simulated related samples with two Han Chinese populations (CHB and CHS) from 1000 Genomes project (Auton et al., 2015) as founders. The simulated kinships have a wide range, including parent offspring, full sibling, first-degree, second-degree, and third-degree cousins. We selected 1.7 million bi-allelic common SNPs to infer kinship (more details in Supplementary). Figure 1 compared performance of different methods (numerical comparisons can be found in Supplementary Table S1 and S2), and we make following observations: 1) The classical method scGRM had noticeable bias in all simulations; 2) popkin (Ochoa and Storey, 2021) appeared to over-corrected and biased upwards, while UKin worked well in correcting the bias; 3) KING performed similarly well with UKin, with UKin performed only slightly better, presumably because KING was benefited from the large number of SNPs we used here compared to the simulations in (Jiang et al., 2022); 4) Kindred and UKin performed similarly well in mean estimates under the alternatives, with Kindred produced slightly better results (mean of the inferred kinships are closest to the theoretical value) in simulations of more distant relationship (cousins); 5) Kindred had the smallest variations under the null, which were the majority of the kinship estimates; 6) Kindred and popkin were the only two methods that produced non-negative estimates on all kinship estimates.

**Figure 1:**
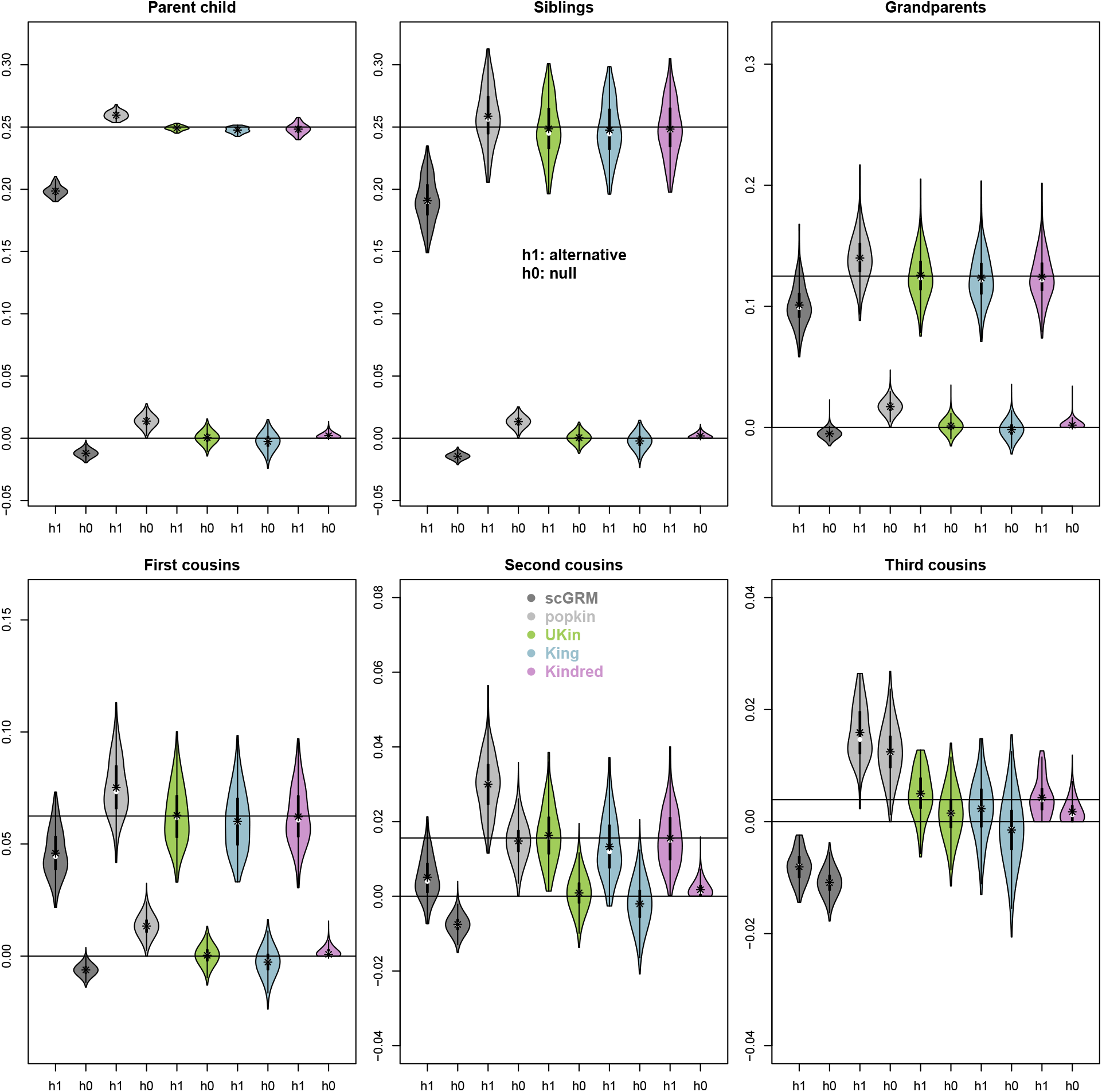
Comparison of kinship estimates by different methods. Results of scGRM (dark gray) were obtained with PLINK. Results of popkin (light gray) were obtained with its R package. Results of UKin (green) were obtained from reimplementation in software Kindred (to take advantage of its multi-threading capacity). Results of King (blue) were obtained from its software King. Results of Kindread (plum) were obtained from its software Kindred. Theoretical values and zero values were marked by horizontal lines and were also given in the table. For each method we showed two violin plots: under the alternative (h1) and under the null (h0). The mean was marked by ⋆. Supplementary Table S1 contains numeric comparisons with mean ± sd under h1, and the percent of negative estimates under h0.

Table 1 was obtained using Han Chinese founders from 1000 Genomes project. To investigate whether Kindred performs similarly well in other founding populations, we conducted additional simulations using diverse founders. Supplementary Figure S3, S4, and S5 compared the deviation between inferred and the truth (instead of the expectation from the pedigree used in Figure 1) for African, East Asian, and European founders respectively. Kindred outcompeted other methods, particularly for harder problems such as kinship between third cousins.

### 3.2 Kinship between admixed samples

Inferring kinship between admixed samples is a difficult problem. The difficulty lies in estimating appropriate allele frequencies. Thornton et al. (2012) solved this problem by using individual-specific allele frequencies, which are admixture proportion-weighted ancestral population frequencies. Here we propose to use a subset of SNPs that have similar allele frequencies between continental populations. If the continental population were taken as the homogenous reference population for IBD for non-admixed samples, then for admixed samples the reference population for IBD has to be the ancestral population predates continental population divergence. This ancestral population can be partially mimicked by selecting a set of SNPs whose allele frequencies are similar across different continental populations, which we call SNPs of small population divergence (SPD). Among 12 million bi-allelic SNPs with minimum 50 minor allele counts (out of total 2504 diplotypes) in the 1000 Genomes project, there are 1.2 million SPD SNPs (details in Supplementary), and we used these SNPs to compute kinship for simulated admixed samples. We also randomly selected common bi-allelic SNPs of 1.2 million, and used these to compute kinship for comparison. We simulated related admixed samples in the same manner as simulating non-admixed samples, the only difference was to choose founders from multiple continents. We chose CEU, YRI, CHB, and CHS as founders to simulate related admixed samples, for these populations show a small extent of inbreeding and a low level of pairwise kinships. Figure 2 demonstrates that our strategy of using SPD SNPs worked well for admixed samples. While randomly selected SNPs produced kinship estimates with much larger variation and the results were perhaps only useful for the first degree relatedness, the selected SPD SNPs produced kinship estimates that were comparable to estimates for non-admixed samples.

**Figure 2:**
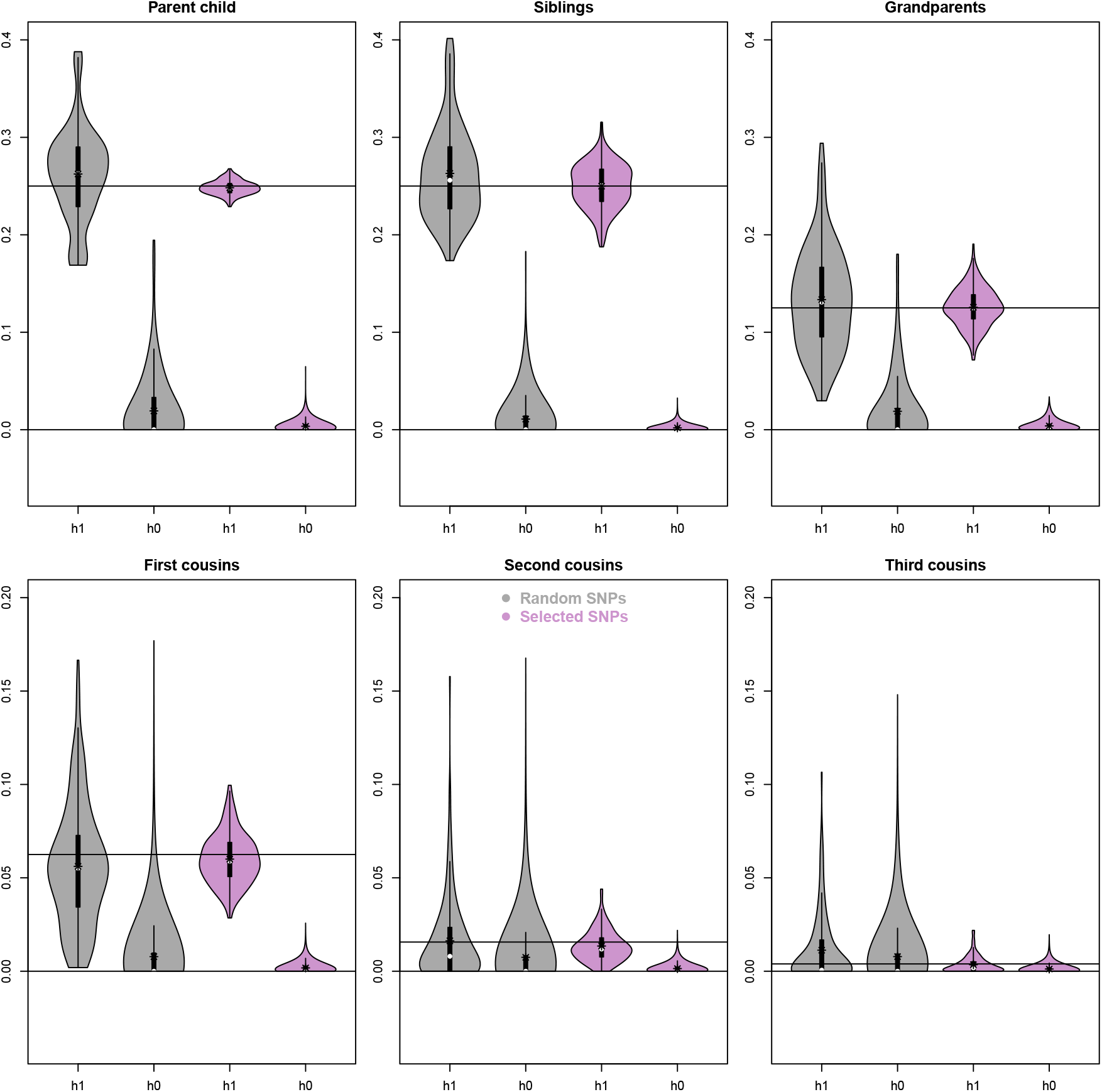
Kindred is effective for admixed samples using SNPs of small population divergence (color plum). Kinship estimated using randomly selected common SNPs (colored in gray) tend to be biased and with large variation. In each panel, theoretical values and zero values were marked by horizontal lines.

### 3.3 Genomic control

One important application of kinship estimates is to control for population stratification and (cryptic) relatedness in genome-wide association studies, either via controlling for top principal components (Price et al., 2006), or incorporate kinship into test statistics (Thornton and McPeek, 2010), or via linear mixed model (Kang et al., 2010; Zhou and Stephens, 2012; Chen et al., 2016). The cohorts in the Framingham Heart Study (FHS), funded by the National Heart, Lung, and Blood Institute (NHLBI), consists many independent three generational pedigrees, nuclear families, trios, duos, and singletons (Kannel et al., 1979). This is a setting in which linear mixed model is effective in controlling for relatedness in the samples. We analyzed 5757 samples with whole genome sequencing data through NHLBI’s TOPMed program (Taliun et al., 2021) and who also have protein immunoassays obtained through the NHLBI’s Systems Approach to Biomarker Research in Cardiovascular Disease (SABRe CVD) Initiative (Ho et al., 2018). Here we analyzed eight proteins that show substantial inflated genomic control values. In Table 2 we compared different kinship estimates in their ability to reduce inflation of test statistics due to the relatedness. All methods are effective in reducing *λ* to near 1, suggesting biases in kinship estimates were well tolerated in the linear mixed model. If measured by deviation from 1, scGRM and King are tied to be the second best, and popkin, UKin and Kindred are tied to be the best. Noticeably, Kindred has the largest mean of *λ* among all methods and is the only one whose mean of *λ* slightly larger than 1.

**Table 2:** Comparison of kinship estimates by their effects on genomic control *λ* values . The single SNP test (to derive genomic control values) using linear mixed model was done with GEMMA. The deviation for each column are calculated from mean of (|*λ −* 1|).

| $\lambda$ | None | scGRM | popkin | UKin | King | Kindred |
| --- | --- | --- | --- | --- | --- | --- |
| lpa | 1.377 | 0.985 | 0.994 | 0.995 | 0.984 | 1.017 |
| pon1 | 1.350 | 0.978 | 0.985 | 0.984 | 0.975 | 1.001 |
| MPO | 1.287 | 0.997 | 0.994 | 0.996 | 1.002 | 1.007 |
| resistin | 1.278 | 0.997 | 0.998 | 0.998 | 1.001 | 1.011 |
| srage | 1.262 | 0.997 | 1.000 | 1.000 | 1.004 | 0.999 |
| cd56 | 1.260 | 0.988 | 0.994 | 0.993 | 0.993 | 1.003 |
| cntn1 | 1.207 | 0.994 | 0.999 | 0.998 | 0.998 | 1.002 |
| CD5L | 1.190 | 1.001 | 1.003 | 1.003 | 1.009 | 1.003 |
| Deviation | 0.276 | 0.008 | 0.005 | 0.005 | 0.008 | 0.005 |
| Mean | 1.276 | 0.992 | 0.995 | 0.995 | 0.996 | 1.005 |

### 3.4 Heritability of height

Using the height data from (Yang et al., 2010), we estimated phenotype variation explained (PVE) by different kinship matrices, where PVE estimates by GCTA were confirmed by GEMMA. The Kindred kinship matrix produced the highest PVE estimates (Table 3). To investigate whether the higher PVE estimates was due to chance or not, we resort to down-sampling. But for the down sampled datasets GCTA produced untenable results due to small sample sizes. With the understanding that Bayesian method performs better with small sample sizes, we reframed the linear mixed model as Bayesian linear regression with a specific prior. Within this framework, we can estimate PVE by seeking MAP (maximum a posterior) estimate of a hyperparameter *η*, which determines the (relative) size of the random effect (details in Appendix). Our down-sampling study showed that Kindred indeed outperformed scGRM (and others) in estimating PVE of the height data, and the difference over 100 down-sampling was statistically significant (for 90% resampling rank test *P* = 4 *×* 10^*−*6^; for 70% resampling rank test *P* = 4 *×* 10^*−*5^; and for 50% resampling rank test *P* = 0.02). It remains to be determined whether this gain in PVE can be translated to power gain in genetic association studies, particularly in multi-omics phenotypes such as proteomics and gene expression assay.

**Table 3:** Mean (*µ*) and standard deviation (*σ*) of heritability estimates of height data and its resampling. The top part of the table was obtained from a single dataset of 3925 samples, where PVE were estimated by GCTA and GEMMA respectively for different kinship estimates; the bottom part of the table was obtained from the resampling study, where 90%, 70% and 50% of the 3925 samples were sampled without replacement in each trial. For each percentage, 100 resampling trials were run and PVEs were estimated using our Bayesian MAP estimates, and mean and standard deviation were computed from the 100 estimates.

|  |  | scGRM | popkin | UKin | King | Kindred |
| --- | --- | --- | --- | --- | --- | --- |
| GCTA | $\mu$ | 0.449 | 0.450 | 0.436 | 0.376 | 0.474 |
| | $\sigma$ | 0.084 | 0.077 | 0.078 | 0.072 | 0.085 |
| GEMMA | $\mu$ | 0.446 | 0.445 | 0.426 | 0.393 | 0.473 |
| | $\sigma$ | 0.084 | 0.077 | 0.079 | 0.071 | 0.085 |
| Resample 90% | $\mu$ | 0.455 | 0.454 | 0.445 | 0.410 | 0.486 |
| | $\sigma$ | 0.045 | 0.033 | 0.038 | 0.032 | 0.041 |
| Resample 70% | $\mu$ | 0.448 | 0.473 | 0.470 | 0.406 | 0.500 |
| | $\sigma$ | 0.086 | 0.081 | 0.086 | 0.078 | 0.088 |
| Resample 50% | $\mu$ | 0.477 | 0.452 | 0.486 | 0.395 | 0.533 |
| | $\sigma$ | 0.151 | 0.137 | 0.111 | 0.126 | 0.160 |

## 4 Discussion

We developed a latent IBD state model to infer kinship and inbreeding coefficients. Our method explicitly models inbreeding when inferring kinships between samples. Kindred makes use of the non-negative least squares method (Lawson and Hanson, 1974) for model fitting, so that our kinship estimates are non-negative, which overcomes difficulties of producing negative estimates that other methods have. In order to use least squares method to fit the latent state model, a unit observation of joint genotypes at a SNP was expanded as a frequency vector, albeit trivial, with one entry as 1 (depending on joint genotypes) and other entries as 0. Compared to a multinomial maximum likelihood approach, the least squares approach we chose here is efficient in computation (Supplementary Table S3), particularly with the readily available modern non-negative least squares method (Bro and De Jong, 1997). The model fitting is quite interesting and we would like to make a brief comment here.

It can be verified that there are two linear dependence in **S**_*p*_ (Table 1). One is Σ_2_ + 2Σ_8_ = Σ_4_ + Σ_6_ + Σ_7_ and the other is *pq*(Σ_1_ + Σ_2_ *−* 2Σ_3_ *−* 2Σ_5_ + 2Σ_7_) = Σ_7_ *−* 2Σ_8_ + Σ_9_. But since the second linear dependence is a function of allele frequency, it disappears when SNPs of different allele frequency bins were used in calculation. Nevertheless, by virtual of the first dependence, the solution to the system 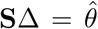 is ot unique. Let **S**^+^ be Moore-Penrose inverse of **S**, then 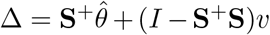 for any vector *v* (Penrose, 1955). Denote *C* = (*I −* **S**^+^**S**)*v*, it can be verified that

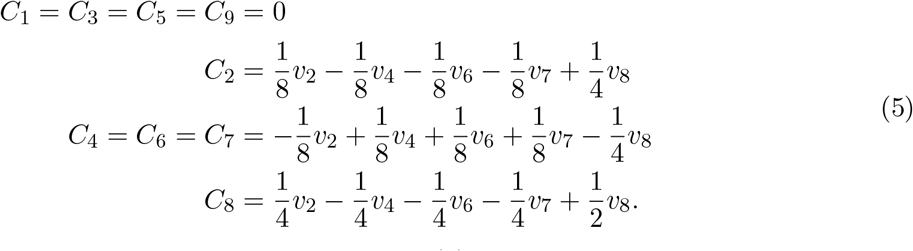

We make the following observations based on Equation (5). First, ∆_1_, ∆_3_, ∆_5_, and ∆_9_ are not affected by *v* and these components have unique solutions. Second, *C*_2_ + *C*_4_ = 0, *C*_2_ + *C*_6_ = 0 and 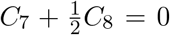, which means, although ∆_2_, ∆_4_, ∆_6_, ∆_7_, and ∆_8_ have infinite many solutions, ∆_2_ + ∆_4_, ∆_2_ + ∆_6_, and 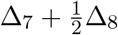 however are invariant. Third, as a consequence to the first two observations, *ϕ* in Equation (1) and *F*_1_ and *F*_2_ in Equation (2) are unique. Csűrös (2014) also observed different coefficients (∆) can generate the same genotype distribution at bi-allelic loci, and made an observation that both kinship and inbreeding coefficients estimates are unaffected by the non-identifiability.

The latent state model can be used as a theoretical framework to analyze other methods such as scGRM. Let *X* and *Y* denote allele counts of genotypes *G*1 and *G*2 in Table 1, *p* is allele frequency of *A* allele, and consider

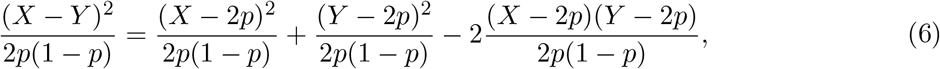

where 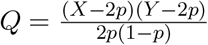 is the quantity calculated in scGRM. The expection of the left hand side in Equation (6) can be directly computed in light of Table 1, and we get *E*(*LHS*) = 4∆_2_ +∆_3_ +3∆_4_ + ∆_5_ + 3∆_6_ +∆_8_ + 2∆_9_. Identifying the expectation of the right hand side is 1 + *F*_1_ + 1 + *F*_2_ *−* 2*Q* and plugging *F*_1_ and *F*_2_ in Equations (2) to get *E*(*RHS*) = 2 + 2∆_1_ + 2∆_2_ + ∆_3_ + ∆_4_ + ∆_5_ + ∆_6_ *−* 2*Q* (more details in Supplementary). Equating the expectations of LHS and RHS and making use the identity 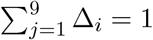, to get the expectation of *Q* as 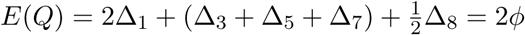, as defined in Equation (1).

A natural question is if scGRM produces unbiased estimates of kinship according to the latent state model framework, then why does it have such noticeable bias? One contributing factor is that the sample allele frequencies were biased from the reference population. Ochoa and Storey (2021) attempted to correct the bias by recalibrating based on the smallest estimates, but our simulations suggested popkin tends to over-correct. On the other hand, Jiang et al. (2022) provided an effective way (UKin) to correct the bias. From above analysis we know *Q* = (1 + *F*_1_ + 1 + *F*_2_ *−LHS*)*/*2, and not accounting for inbreeding effectively sets *F*_1_ = *F*_2_ = 0, which reduces *Q* and thus contributes to excessive negative entries in scGRM. From the perspective of latent state model, however, not modeling inbreeding is effectively setting loading probabilities of the first six latent states as zero, and those excessive allele sharing caused by inbreeding are more likely to be assigned to latent states Σ_7_ and Σ_8_ than Σ_9_ (Figure 1), resulting in slight upward bias (the amount of bias were limited by the kinship between founders). Supplementary Figure S6 demonstrated this intuition via simulations.

The definitions of inbreeding and kinship hinge on IBD, while IBD is defined relative to a reference population, where different alleles in that reference population are considered *not* IBD (Wang, 2016; Goudet et al., 2018). The defining feature of a reference population is its marginal allele frequencies. (Arguably, the best feature of a reference population is its haplotype frequencies, but that is difficult to obtain and not easy to work with.) Both scGRM and UKin used allele frequencies estimated from samples, which implicitly assumed samples were taken from reference populations and allele frequencies estimates were unbiased. Both were strong assumptions. It has been pointed out that one has to take into account of pedigree or cryptic relatedness to obtain unbiased estimates of allele frequencies (Sara McPeek et al., 2004). So an iterative procedure to refine estimates of allele frequencies and kinship over and over might help to alleviate bias and negativeness observed in these methods. Kindred allows users to specify allele frequencies through a tag in VCF files, which effectively allows users to specify a reference population.

Although mis-specifying a reference population will bias the kinship (and inbreeding coefficient) estimates, there in fact has nothing wrong with such a practice if the results were interpreted accordingly. For example, when using Africans as reference population, kinship estimates between East Asians were elevated compared to using East Asian as reference population. Since African alleles are more diverse than East Asian alleles, it is only reasonable that East Asians are modestly “inbred” with reference to the African alleles. In fact, through lens of the African alleles, certain patterns in East Asian samples become more apparent. Figure 3 showed such an example. Pairwise PC plots formed three distinct clusters (between PC2 and PC3) when using African allele frequencies to compute kinship matrix on chromosome 17 (Figure 3A upper triangle). Such clustering is visible but has no distinct separation from PC plots obtained from East Asian allele frequencies (Figure 3A lower triangle), and there is no visible clustering in PCs from scGRM (Figure 3D lower triangle), although the coloring suggested samples of the same color indeed aggregate (Figure 3D upper triangle). In other words, Figure 3A upper triangular plots makes the discovery possible. The clustering was not caused by population structure in East Asians (Supplementary Figure S7). We used a linear combination of PC2 and PC3 to derive a phenotype that cleanly separate samples into three clusters (Figure 3B), and performed single SNP test using BIMBAM (Servin and Stephens, 2007). Interestingly, SNPs strongly associated with the derived phenotype clustered around centromere region (Figure 3C). We used PC3 for scGRM as a derived phenotype (Figure 3E) and performed single SNP test again and observed similar pattern but with reduced magnitude in Bayes factors (Figure 3F). A recent study suggested that haplotype spanning centromere regions may be introgressed from archaic DNA (Langley et al., 2019), which might explain why using African alleles as reference we obtained more distinct clusters in East Asians on chromosome 17.

**Figure 3:**
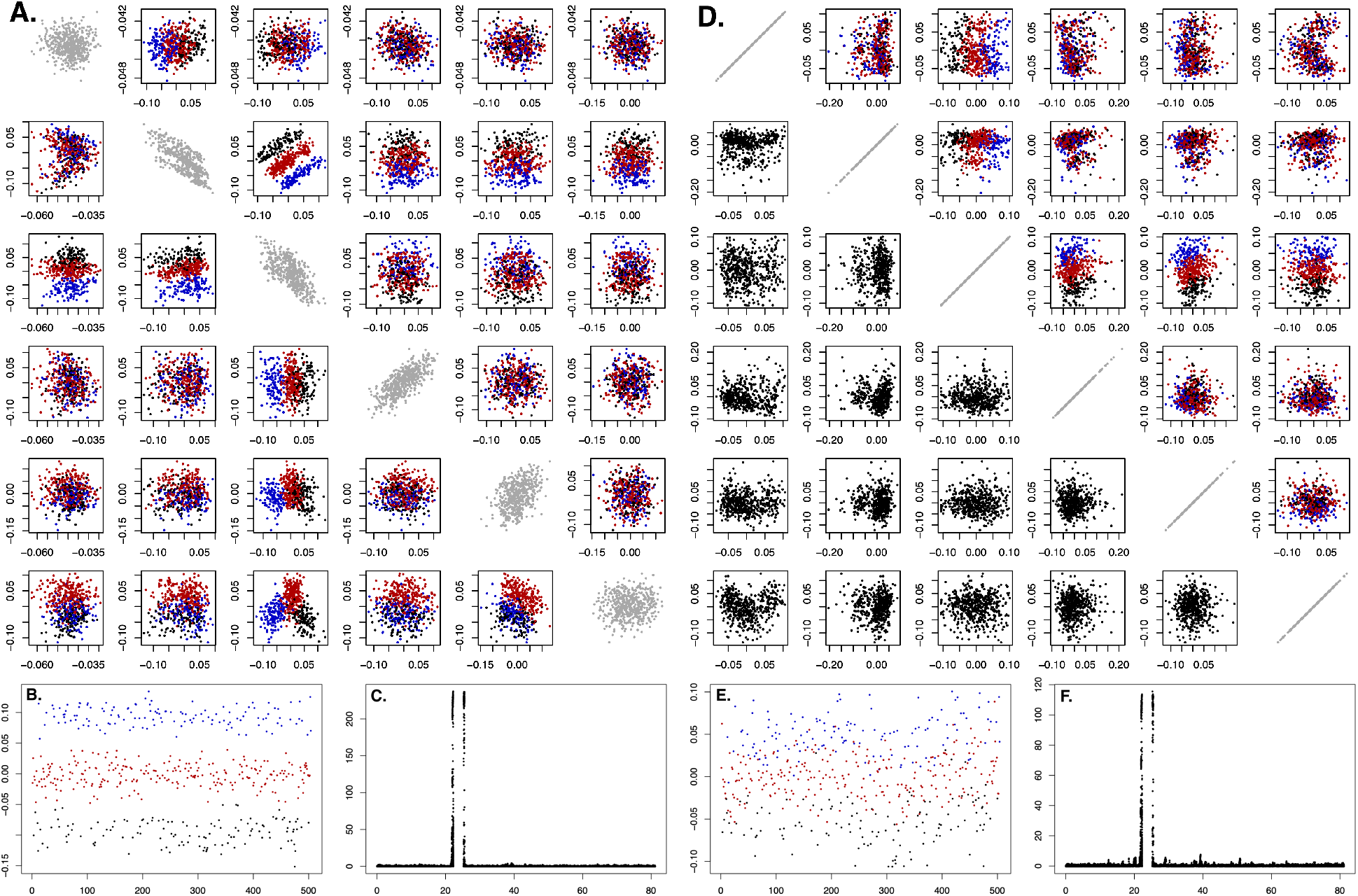
East Asian samples clustering pattern on Chr17. Panel A: pairwise plot of top six Kindred PCs. The lower triangular plots were PCs of kinship matrix inferred with East Asian allele frequencies. The upper triangular plots, African allele frequencies. The diagonals are plots of *j*-th PC from one vs *j*-th PC from the other. Panel B: phenotypes derived from PC2 and PC3 in upper triangular plots in Panel A. Individuals were assigned into three clusters by Kmeans method and marked in three different colors. (Samples used the same color in Panels A,B,D,E.) Panel C: Manhattan plot of single SNP (log_10_) Bayes factors between common biallelic SNPs on Chr17 and derived phenotypes in Panel B. Panel D: pairwise plots of top six scGRM PCs. Without coloring, the relevant groups unidentifiable (lower triangle plots). In colored samples (upper triangle), three groups of samples aggregate, but not form distinct clusters as in Kindred PCs. Panel E: derived phenotype from PC3 in Panel D. Panel F: Manhattan plot of single SNP (log_10_) Bayes factors between common biallelic SNPs on Chr17 and derived phenotypes in Panel E. Note the y-axis range in Panel F is half of that in Panel C.

Using a set of SNPs with small population divergence (SPD) to mimic allele frequencies of the ancestral population, Kindred can estimate kinship for admixed samples with high accuracy. (We chose not to use the traditional *F*_*st*_ to the measure the population divergence out of the consideration that *F*_*st*_ emphasize pairwise difference, while we wanted similar allele frequencies in at least three continental populations.) We recommend SPD SNPs to be used when analyzing samples with more than one continental origin. Arguably, when jointly analyzing diverse populations (in addition to analyzing admixed samples), using the SNPs of small population divergence is more sensible than all alternatives. We inferred realized kinship matrices for all 2504 samples in the 1000 Genomes project, using either 1.2 million SPD SNPs or 1.2 million randomly selected common SNPs. Figure 4 compared their top five PCs. PCs from SPD SNPs still cleanly separates continental populations, which suggests that although each individual SNPs are non-informative to ancestry, combined they are distinctly informative to ancestry. For PCs from randomly selected common SNPs, a striking feature is that PC1 was dominated by Africans samples, which have very little variation in other four leading PCs. This feature appears to be unique to Kindred.

**Figure 4:**
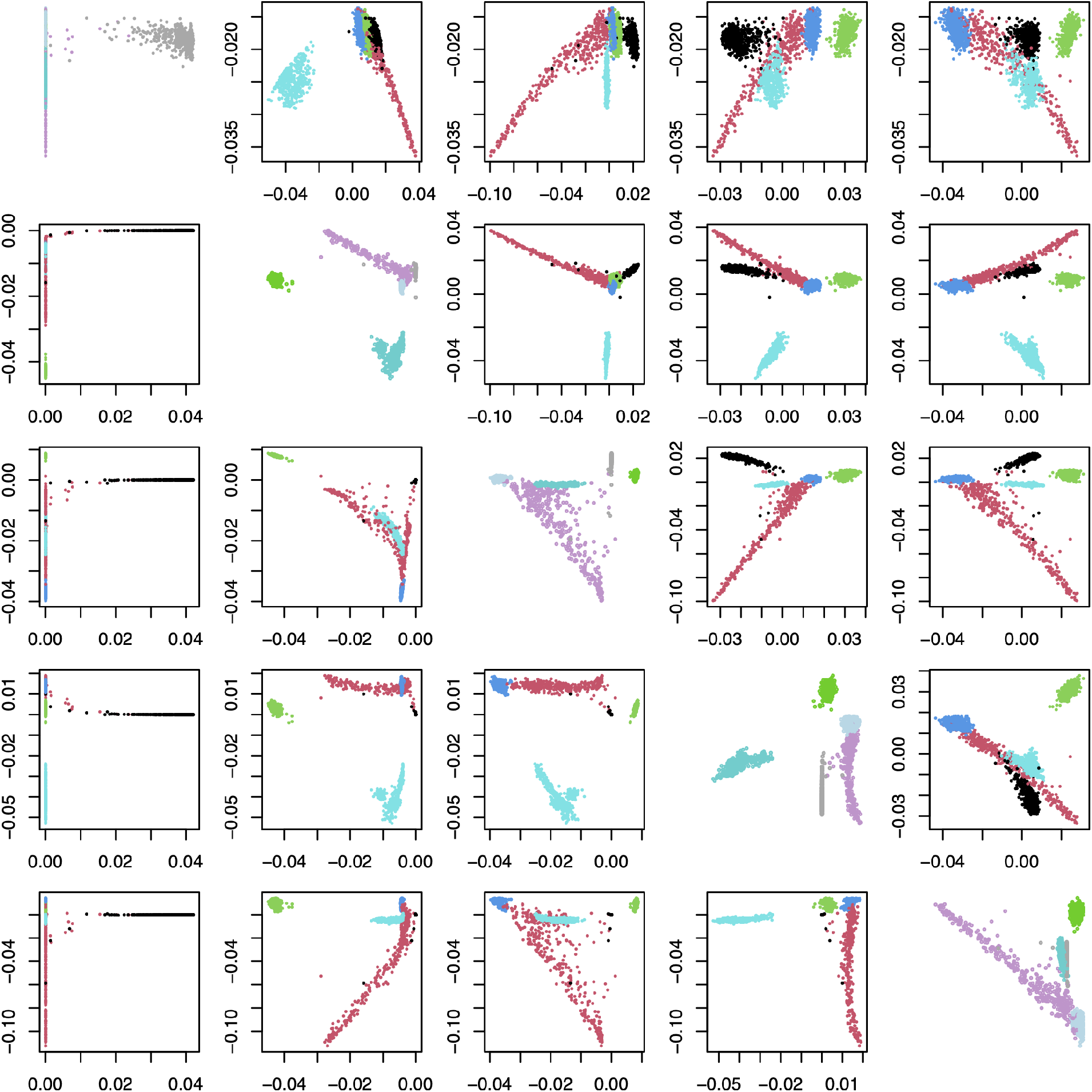
PC plots of 1000 genomes samples. The upper triangle are pairwise plots of top five PCs inferred with small population divergence (SPD) SNPs. The lower triangle, randomly selected common (RSC) SNPs. The diagonals are plots of *j*-th PC from SPD SNPs vs *j*-th PC from RSC SNPs, with *j* = 1, 2, 3, 4, 5. Five continental samples are Africans (in black), Americans (in red), East Asians (in green), Europeans (in blue), and South asians (in cyan). On diagonal their colors are in gray, plum, light green, light blue, and cyan.

## 5 Appendix

### 5.1 Bayes factor of linear mixed model

Consider a linear model

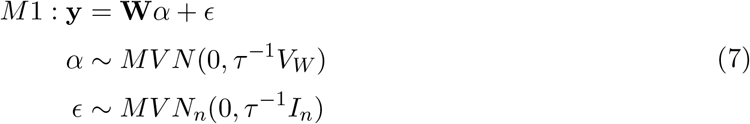

where **W** is a *n × w* representing *w* nuisance covariates. Adding a random effect **u** we have a new model

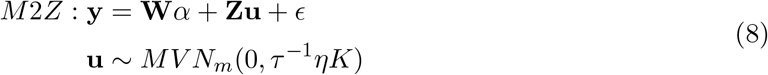

where *Z* is *n×m* matrix presenting the loading, and *K* is *m×m* covariance matrix. Let *Q* and *D* be eigen decomposition such that *ZKZ*^*t*^ = *QDQ*^*t*^, where *D* = *diag*(*d*_1_, …, *d*_*n*_) with *d*_1_ *≥ d*_2_ *≥ · · · ≥ d*_*n*_ and *QQ*^*t*^ = *I*. Equations 8 can be rewritten as

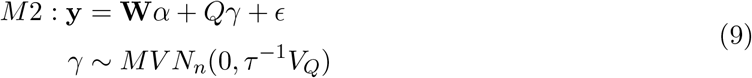

where *V*_*Q*_ = *ηD*. To see this, *E*(*Z***uu**^*t*^*Z*^*t*^) = *ηZKZ*^*t*^ = *ηQDQ*^*t*^ = *E*(*Qγγ*^*t*^*Q*^*t*^).

Since models M1 and M2 are nested, it is understood that the distribution assumption and prior specification used in a simpler model carry over to the more complex model. By specifying a Gamma prior on *τ* we have normal-inverse-gamma on a linear model (c.f. Servin and Stephens, Zhou and Guan).

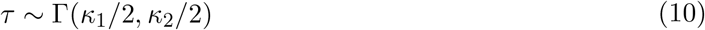

It’s clear that from a Bayesian perspective, a linear mixed model is just a linear model with a specific prior. For *M* 1 after integrating out *α* and *τ* and letting *κ*_1_, *κ*_2_ *→* 0, we have

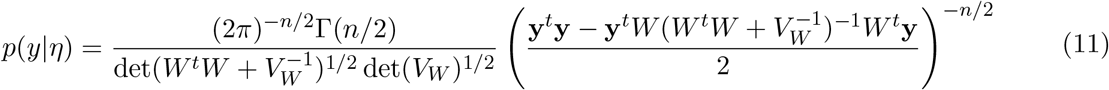

Treating *M*1 as null and *M*2 as alternative, we compute *BF*_21_ in a closed form with the above prior specification. Denote *X* = (*Q, W*) and 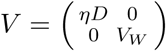, we have

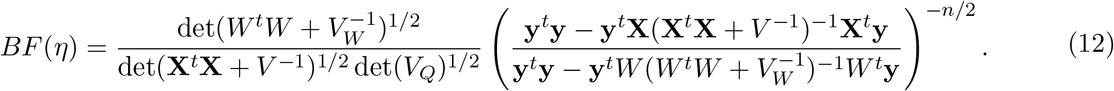

Let *V*_*W*_ *→ ∞*, we have 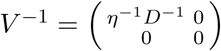 and

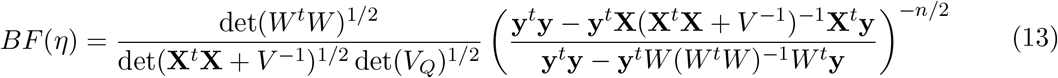

### 5.2 Bayesian estimates of PVE

Bayes factor (13) can be evaluated efficiently for different *η*. As 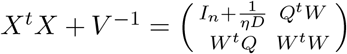, we compute its determinant using the identity

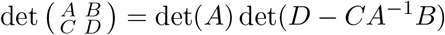

to get

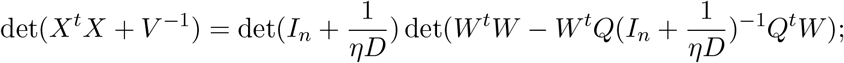

and its inverse using the identities

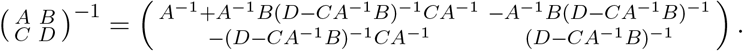

Denote 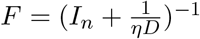, *M* = (*W* ^*t*^*W − W* ^*t*^*QFQ*^*t*^*W*)^*−*1^ to get

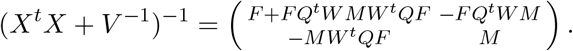

Note these computations only involve inexpensive matrix multiplying vector, without expensive matrix multiplication and matrix inversion. The only expensive calculation is the eigendecomposition to obtain Q and D, which only needs to be done once. A R script to compute Bayes factor (Equation 13) is available in the Supplementary.

With efficient evaluation of BF at hand, we can find 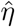 that maximize *BF* (*η*) by Nelder-Meade algorithm, and this is a MAP (maximum a posterior) estimator. With 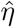 we can compute 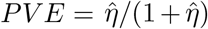. In Supplementary Figure S8, we demonstrate two properties of the PVE via simulations: 1) Bayesian estimates recovered the true PVE; and 2) Bayesian estimates are consistent with those GCTA estimates.

## 6 Acknowledgements

This research is supported by the Division of Intramural Research of the National Heart, Lung, and Blood Institute, Bethesda, MD (D.L. Principal Investigator). We thank Nick Martin and the Queensland Institute of Medical Research for making the Australia height data available to us.

